# Donor-Aware Reanalysis of Neuronal Nuclei Supports STMN2 Depletion Associated with Nuclear TDP-43 Loss in FTD and ALS

**DOI:** 10.64898/2026.10.27.762172

**Authors:** Abdullah Chaudhary, Saiqi Yao

## Abstract

**Background:** Nuclear TDP-43 depletion is a central pathological feature of amyotrophic lateral sclerosis (ALS) and frontotemporal dementia (FTD), but donor-to-donor variation can obscure transcriptomic associations in small postmortem cohorts. We reanalyzed public RNA-seq data from FACS-isolated NeuN-positive frontal-cortex neuronal nuclei to quantify gene-expression changes associated with nuclear TDP-43 loss while explicitly accounting for donor structure.

**Methods:** The supplied GSE126542-derived matrix contained 62,754 genes across 12 libraries from seven donors, including five complete matched pairs. Source labels “TDP-43 positive” and “TDP-43 negative” were harmonized as nuclear TDP-43 retained and lost, respectively. After validation against featureCounts-assigned read counts, 44,225 genes with at least 10 total counts were analyzed using PyDESeq2. The primary model was ∼ donor + nuclear_status; status-only and diagnosis-adjusted models were sensitivity analyses. The contrast was TDP-43 loss minus retained.

**Results:** The donor-aware model identified 3,442 genes at false discovery rate (FDR) < 0.05: 1,865 lower and 1,577 higher in TDP-43-loss nuclei. STMN2 was consistently depleted (log2 fold change = −1.813, 95% Wald interval −2.229 to −1.396; FDR = 6.22 × 10^−15^), corresponding to approximately 71.5% lower expression. Normalized STMN2 counts were lower in TDP-43-loss nuclei in all five complete donor pairs. TARDBP expression was also lower (log2 fold change = −0.606; FDR = 4.98 × 10^−6^). In contrast, UNC13A whole-gene expression was modestly higher in the donor-aware model (log2 fold change = 0.372; FDR = 0.00622) and was not significant in either sensitivity model.

**Conclusions:** Donor-aware analysis supports a robust association between nuclear TDP-43 loss and STMN2 depletion in this neuronal-nuclei cohort, alongside extensive but model-dependent transcriptomic variation. Whole-gene UNC13A expression cannot be used to infer UNC13A cryptic-exon inclusion or functional UNC13A depletion. No splice-junction, percent-spliced-in, isoform, or protein measurements were available; mechanistic claims therefore require junction-level and experimental validation.

## Introduction

TAR DNA-binding protein 43 (TDP-43) is an RNA-binding protein whose cytoplasmic mislocalization and aggregation are pathological hallmarks of ALS and a major subset of FTD [1]. In addition to the consequences of cytoplasmic aggregation, loss of nuclear TDP-43 function disrupts RNA processing. A well-established function of nuclear TDP-43 is repression of nonconserved cryptic exons; when this repression fails, otherwise excluded sequences can enter mature transcripts [2]. This loss-of-function framework links a pathological observation—nuclear TDP-43 depletion—to specific RNA-processing outcomes, but it also creates a measurement requirement: gene-level counts and exon-inclusion measurements are not interchangeable.

Two genes illustrate why that distinction matters. STMN2 encodes stathmin-2, a regulator of microtubule dynamics and axonal growth and repair. Experimental studies have connected TDP-43 dysfunction to altered STMN2 RNA processing and reduced stathmin-2 output [3,4]. UNC13A has likewise emerged as a key TDP-43-sensitive gene, with cryptic-exon inclusion, nonsense-mediated decay, and genetic variation jointly shaping its interpretation [5,6]. Mis-spliced transcripts can also generate de novo protein species, emphasizing that transcript structure and protein consequences require direct assays [7]. A whole-gene count matrix alone cannot resolve any of these event-level mechanisms.

The public GSE126542 series generated by Liu et al. provides an unusual opportunity to study nuclear TDP-43 loss in disease-relevant cells [8]. Rather than profiling only unsorted cortex, the source study used subcellular fractionation and FACS enrichment to isolate NeuN-positive neuronal nuclei with or without detectable nuclear TDP-43. Matched nuclei from the same donor make a donor-aware analysis possible, reducing the influence of stable interindividual differences. The supplied reanalysis matrix contains 12 such libraries from seven donors, five of whom have both nuclear states represented.

We therefore performed a secondary, count-level reanalysis with three aims: (i) verify the supplied matrix and sample annotations; (ii) estimate global and prespecified marker-gene associations using a primary donor-aware model and two sensitivity models; and (iii) define precisely what the data can and cannot establish about STMN2, UNC13A, and cryptic-exon biology. We hypothesized that donor-aware modeling would preserve the expected STMN2 association while revealing heterogeneity that is obscured by treating libraries as independent. We did not attempt to infer splicing, pathway activation, or protein-level consequences from gene-level counts.

## Methods

### Study design and source data

This study is a secondary analysis of publicly available, deidentified RNA-seq data from GSE126542 (BioProject PRJNA522295), originally reported by Liu et al. [8]. Source metadata identify frontal-cortex, NeuN-positive neuronal nuclei sequenced as paired-end libraries on an Illumina HiSeq 2500. The source labels “TDP-43 positive” and “TDP-43 negative” denote retained and lost nuclear TDP-43, respectively. Throughout this manuscript, “loss” refers to the source nuclear-TDP-43 annotation rather than a new immunohistochemical measurement. Source metadata annotate a C9orf72 hexanucleotide repeat-expansion background; this restricts the populations to which the findings can be generalized.

The full series contains 16 RNA-seq libraries. Two unsorted libraries (SRR8571943 and SRR8571946) were outside the neuronal-nuclei comparison. Fourteen libraries were eligible by cell-type and TDP-43 annotations, but the supplied matrix contained 12. The rationale for omitting the two otherwise eligible sorted libraries, SRR8571945 and SRR8571947, was not available in the supplied materials. The analyzed cohort therefore comprised six TDP-43-retained and six TDP-43-loss libraries from seven donors, with five complete pairs and two singleton donors (Table 1; Fig. 1).

**Table 1.** Donor-level structure of the analyzed neuronal-nuclei cohort.

| Donor | Diagnosis | Sex | Retained library | Loss library | Pairing status |
|---|---|---|---|---|---|
| S1 | FTD | Female | SRR8571937 | SRR8571938 | Complete pair |
| S2 | FTD–ALS | Male | SRR8571939 | SRR8571940 | Complete pair |
| S3 | FTD | Female | SRR8571941 | SRR8571942 | Complete pair |
| S4 | FTD | Female | SRR8571944 | — | Retained only |
| S5 | FTD | Male | — | SRR8571948 | Loss only |
| S6 | FTD–ALS | Male | SRR8571949 | SRR8571950 | Complete pair |
| S7 | FTD–ALS | Male | SRR8571951 | SRR8571952 | Complete pair |

All libraries were frontal-cortex, FACS-isolated NeuN-positive neuronal nuclei. S4 and S5 each have an eligible counterpart in the full series that is absent from the supplied matrix. No healthy-control libraries were present.

**Figure 1.**
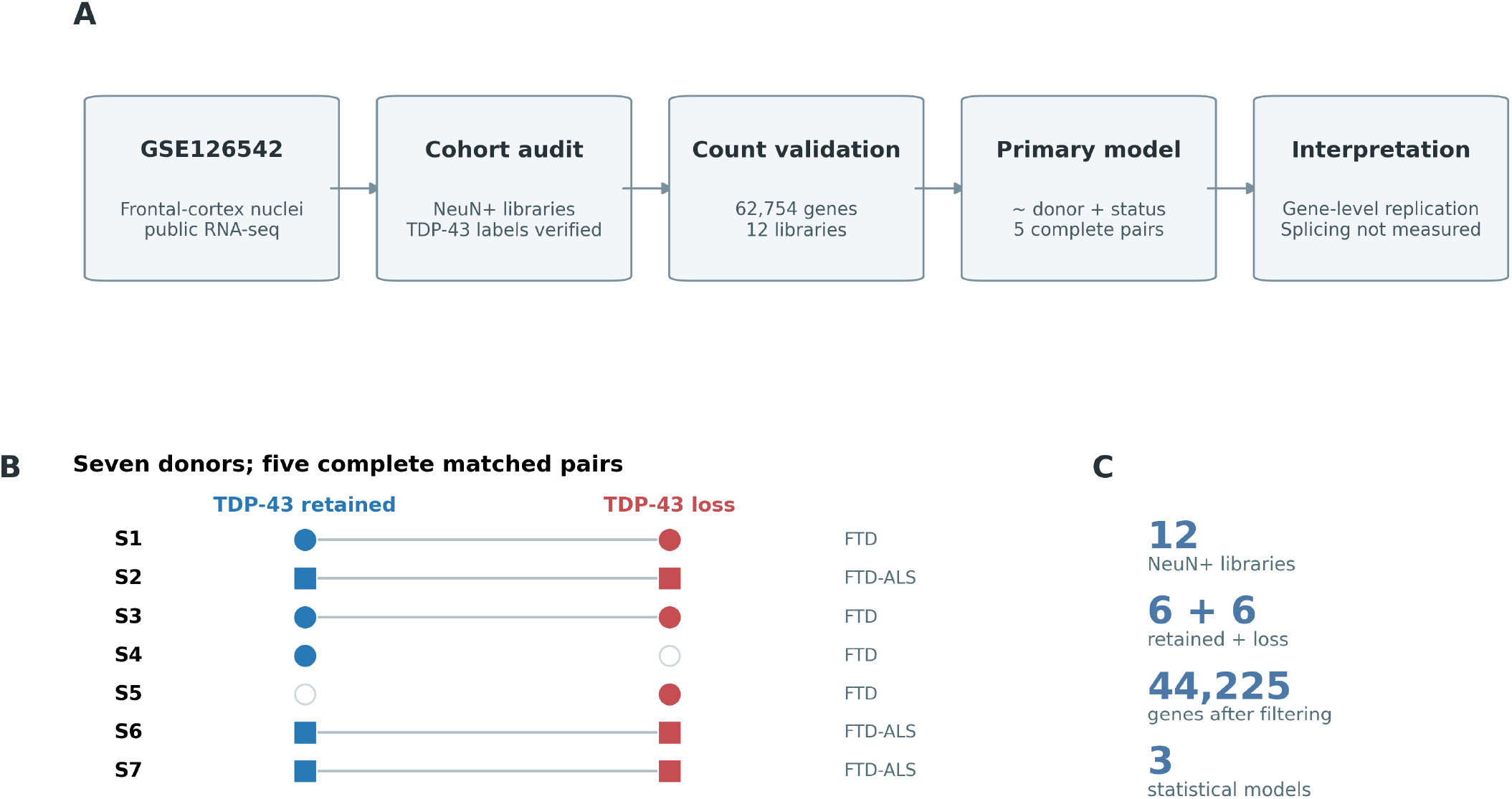
Study design and donor structure. (A) Secondary-analysis workflow from public GSE126542 metadata and the supplied count matrix to donor-aware differential expression and gene-level interpretation. (B) Seven donors with nuclear TDP-43-retained and TDP-43-loss libraries; lines connect complete matched pairs, open symbols indicate absent states, circles indicate FTD, and squares indicate FTD–ALS. (C) Key cohort and analysis dimensions. Splicing and PSI were not estimated.

### Metadata harmonization and analysis contrast

Run accessions, GEO sample identifiers, BioSample identifiers, tissue, cell type, diagnosis, sex, library layout, and sequencing instrument were cross-checked against the verified source-metadata table. Donor identifiers S1–S7 were recovered from the source sample titles. The analysis variable nuclear_status was encoded as Positive for retained nuclear TDP-43 and Negative for nuclear TDP-43 loss to match the supplied model code. Every differential-expression contrast was Negative versus Positive—TDP-43 loss minus retained. Consequently, negative log2 fold changes indicate lower expression in TDP-43-loss nuclei.

### Count-matrix validation

The supplied counts_matrix.txt file was read as an unheadered Ensembl-gene-by-library integer matrix, with library order recovered from raw_counts.txt.summary. Validation required unique gene identifiers, no missing values, and nonnegative integer counts. Column sums were compared exactly with featureCounts Assigned counts. The matrix contained 62,754 genes and 12 libraries; 10,564 genes had zero counts in every library. Library sums matched the Assigned counts for all libraries, ranging from 18,584,661 to 29,152,965 assigned reads (median 24,343,171). Genes with at least 10 counts summed across all 12 libraries were retained, yielding 44,225 genes. The featureCounts summary was also used to visualize assigned, multimapping, no-feature, and ambiguous categories; these categories were not interpreted as STAR alignment rates (Fig. S1).

### Reference FASTQ-to-counts workflow and its execution status

A configurable paired-end workflow is supplied as code/01_bulk_rnaseq_pipeline.sh to document how an equivalent FASTQ-to-counts pipeline can be constructed. It uses fasterq-dump for accession retrieval, FastQC for read-quality inspection, Trimmomatic for adapter and quality trimming [9], STAR for splice-aware alignment [10], samtools for BAM indexing, and featureCounts for exon-level assignment aggregated by gene identifier [11]. The script exposes sample accessions, reference FASTA, GTF, genome index, adapter file, thread count, and featureCounts strandedness as configuration inputs.

This FASTQ workflow was not executed for the reported reanalysis. All statistical results begin from the supplied count matrix and its featureCounts summary. Accordingly, we do not report new alignment percentages, duplicate rates, trimming yields, or strandedness checks. The supplied matrix uses Ensembl gene identifiers, but the exact genome build, annotation release, and original alignment settings could not be independently reconstructed from the supplied files; no unverified release number is asserted. GENCODE is cited as the relevant reference-annotation framework [12], not as evidence that a particular release was used.

### Differential-expression models

Differential expression was estimated with PyDESeq2 version 0.5.4 under Python 3.12.12, implementing DESeq2-family negative-binomial modeling [13,14]. The primary design was ∼ donor + nuclear_status. This model estimates the nuclear-status effect while accounting for donor-specific baseline differences; its within-donor status contrast is identified principally by the five complete pairs. The two singleton donors contribute to model fitting but do not themselves supply a matched status comparison. Two sensitivity designs were fitted: ∼ nuclear_status and ∼ diagnosis + nuclear_status.

For each model, DeseqDataSet was fitted with Cook’s-distance refitting enabled. DeseqStats used the contrast [nuclear_status, Negative, Positive], alpha = 0.05, Cook’s-distance filtering, and independent filtering. Result tables contain baseMean, log2 fold change, standard error, Wald statistic, nominal P value, and Benjamini–Hochberg-adjusted P value. Genes with FDR < 0.05 were considered statistically significant. A secondary threshold of FDR < 0.05 and absolute log2 fold change ≥ 1 was used only to describe large-effect subsets, not to redefine the primary significance criterion. Complete result tables are supplied for all three models.

### Descriptive normalization, PCA, and heatmaps

For visualization only, standard median-of-ratios size factors were calculated from the filtered matrix using genes with nonzero counts in every library. Normalized counts were transformed as log2(normalized count + 1). Principal-component analysis (PCA) used the 2,000 genes with the highest variance after transformation, with gene-wise centering. Matched donor pairs were connected in the PCA plot to make within-donor structure visible; PCA was descriptive and was not used for batch correction or significance testing.

The leading-gene heatmap displays 30 protein-coding genes selected by the smallest donor-aware adjusted P values among genes meeting FDR < 0.05, absolute log2 fold change ≥ 1, baseMean ≥ 100, and successful symbol annotation. Log2-normalized expression was converted to row Z scores, and rows were ordered by average-linkage hierarchical clustering using correlation distance. Libraries remained ordered by donor and nuclear status rather than being treated as independent validation clusters. Gene-selection statistics are provided in Table S9.

### Prespecified marker analysis and effect-size summaries

STMN2 (ENSG00000104435), UNC13A (ENSG00000130477), and TARDBP (ENSG00000120948) were prespecified in the analysis script because of their relevance to TDP-43 biology. Their normalized counts were plotted by nuclear status with lines connecting complete donor pairs. Model estimates were summarized as log2 fold change ± 1.96 standard errors to give 95% Wald intervals. Approximate percent change was calculated as 100 × (2^log2 fold change − 1). For descriptive paired review, within-donor STMN2 log2 ratios of normalized counts were calculated for the five complete pairs. These ratios were not substituted for the model-based estimates.

### Gene annotation and derived tables

Ensembl identifiers were annotated through the MyGene.info v3 query service using ensembl.gene scope and human-specific symbol, name, and gene-type fields. Queries covered all donor-aware FDR-significant genes, the 50 leading genes by adjusted P value, and the three markers. The first returned hit per queried identifier was used for display after all raw responses were preserved. Unmapped identifiers were retained and exported rather than silently removed. Annotation was used for labeling and heatmap eligibility, not for differential-expression inference.

### Cryptic-exon evidence boundary

The available inputs contained gene-level counts and featureCounts summaries, not splice-junction counts, junction-aware alignments, transcript-level quantification, or event-level percent-spliced-in (PSI) estimates. We therefore did not calculate cryptic-exon inclusion, differential exon usage, isoform switching, nonsense-mediated decay, or cryptic-peptide abundance. A change in whole-gene expression can coexist with, occur independently of, or fail to detect an event affecting only a subset of transcripts. Figure S3 explicitly separates measured nuclear-status labels and whole-gene expression from unmeasured splicing and protein consequences.

### Reproducibility and statistical reporting

The analysis package separates data, verified metadata, model results, annotation outputs, figure code, and manuscript files. SHA-256 checksums were recorded for the count matrix, featureCounts summary, and verified metadata. The differential-expression script supports validation-only execution and configurable model selection. The figure script regenerates eight figures in PNG, PDF, and TIFF formats and exports PCA coordinates, normalized marker counts, heatmap selection criteria, size factors, and model-concordance tables. All P values reported in the main text are two-sided Wald tests unless otherwise stated. No independent external cohort was analyzed.

## Results

### Cohort structure and count integrity were verified

The analyzed matrix comprised 12 NeuN-positive neuronal-nuclei libraries: six with nuclear TDP-43 retained and six with nuclear TDP-43 loss. These libraries represented seven donors and five complete matched pairs. Donors S4 and S5 contributed only retained and loss libraries, respectively, despite eligible counterparts being present in the full source series (Table 1; Fig. 1). This incomplete pairing and undocumented subset selection were retained as explicit limitations rather than repaired by imputation.

All 62,754 gene identifiers were unique, all counts were nonnegative integers, and no values were missing. Exact agreement between matrix column sums and featureCounts Assigned counts supported internal count integrity. Assigned-read depth varied approximately 1.57-fold across libraries but no library was an extreme low-count outlier (Fig. 2A). After minimum-count filtering, 44,225 genes remained for model fitting.

**Figure 2.**
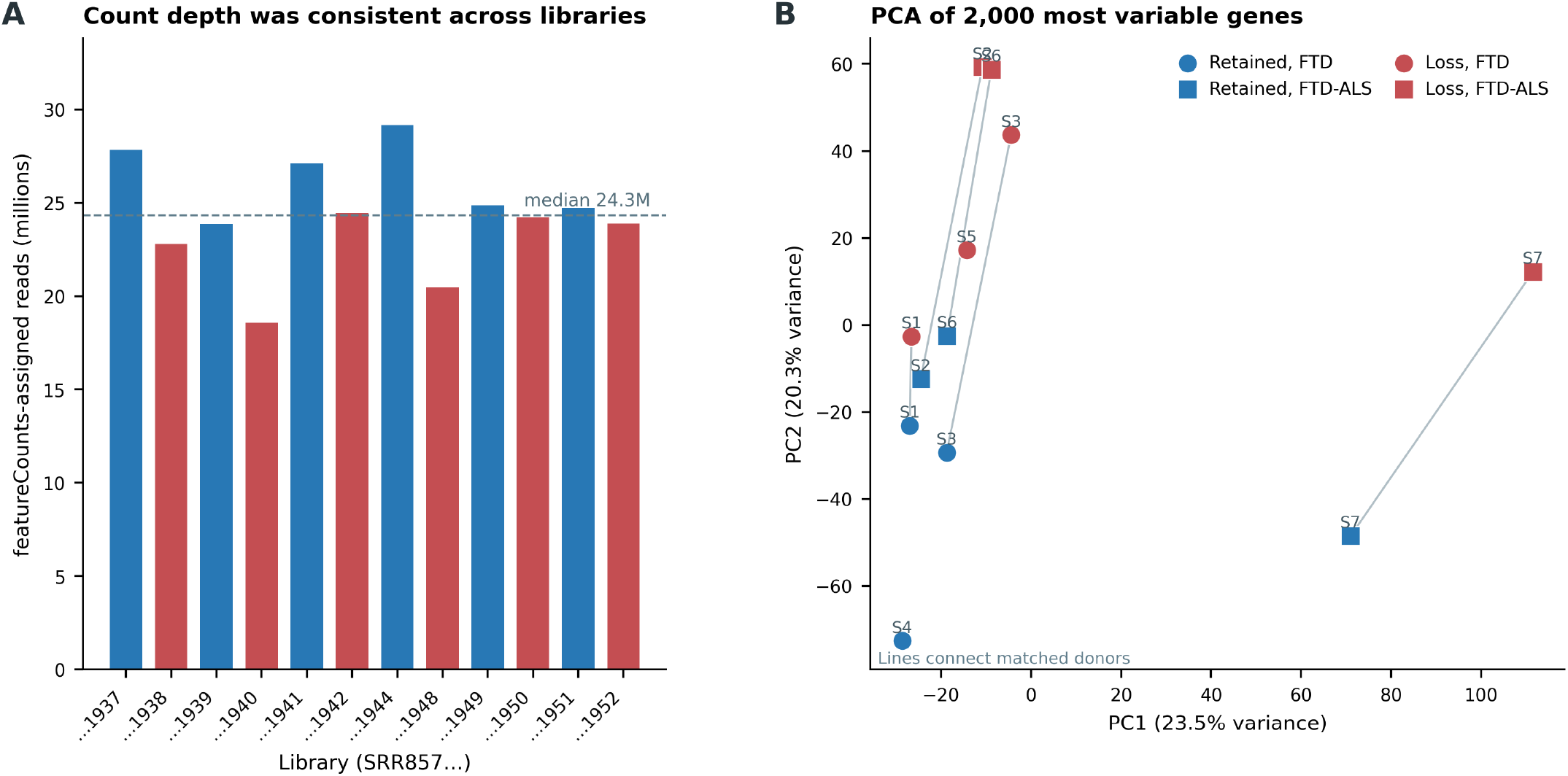
Count depth and global expression structure. (A) featureCounts-assigned reads per library; the dashed line marks the median of 24.3 million assigned reads. Colors indicate nuclear TDP-43 status. (B) PCA of log2(normalized count + 1) values for the 2,000 most variable genes. Lines connect complete matched donor pairs; shapes indicate diagnosis and colors indicate nuclear status. PCA is descriptive and was not used for inference.

PCA of the 2,000 most variable genes placed 23.5% and 20.3% of total variance on PC1 and PC2, respectively (Fig. 2B; Table S6). Neither diagnosis nor nuclear status alone produced a clean global separation. Matched-donor connectors showed heterogeneous within-pair movement, consistent with substantial donor and library variation. Because PCA is unsupervised and dominated by high-variance genes, it was not used to judge the validity of individual marker associations.

### Donor-aware modeling identified widespread but model-dependent expression differences

In the primary donor-aware model, 30,506 genes received adjusted P values and 3,442 met FDR < 0.05. Of these, 1,865 were lower and 1,577 were higher in TDP-43-loss nuclei; 1,528 additionally had absolute log2 fold change ≥ 1 (Fig. 3A,B; Table 2). The status-only model identified 1,295 significant genes (823 lower, 472 higher), and the diagnosis-adjusted model identified 1,495 (944 lower, 551 higher). The larger number of discoveries in the donor-aware model should be interpreted as model-dependent inference in this small cohort, not as an independent replication or a guarantee of greater biological specificity.

**Table 2.** Differential-expression summaries across the three fitted models.

| **Model** | **Design** | **Adjusted P available** | **FDR < 0.05** | **Lower in loss** | **Higher in loss** | **FDR + \|LFC\| ≥ 1** |
|---|---|---|---|---|---|---|
| Status only | ~ nuclear_status | 34,306 | 1,295 | 823 | 472 | 1,027 |
| Diagnosis adjusted | ~ diagnosis + nuclear_status | 34,345 | 1,495 | 944 | 551 | 1,131 |
| Donor aware | ~ donor + nuclear_status | 30,506 | 3,442 | 1,865 | 1,577 | 1,528 |

All contrasts are nuclear TDP-43 loss minus retained. “FDR + \|LFC\| ≥ 1” denotes genes with FDR < 0.05 and absolute log2 fold change at least 1.

**Figure 3.**
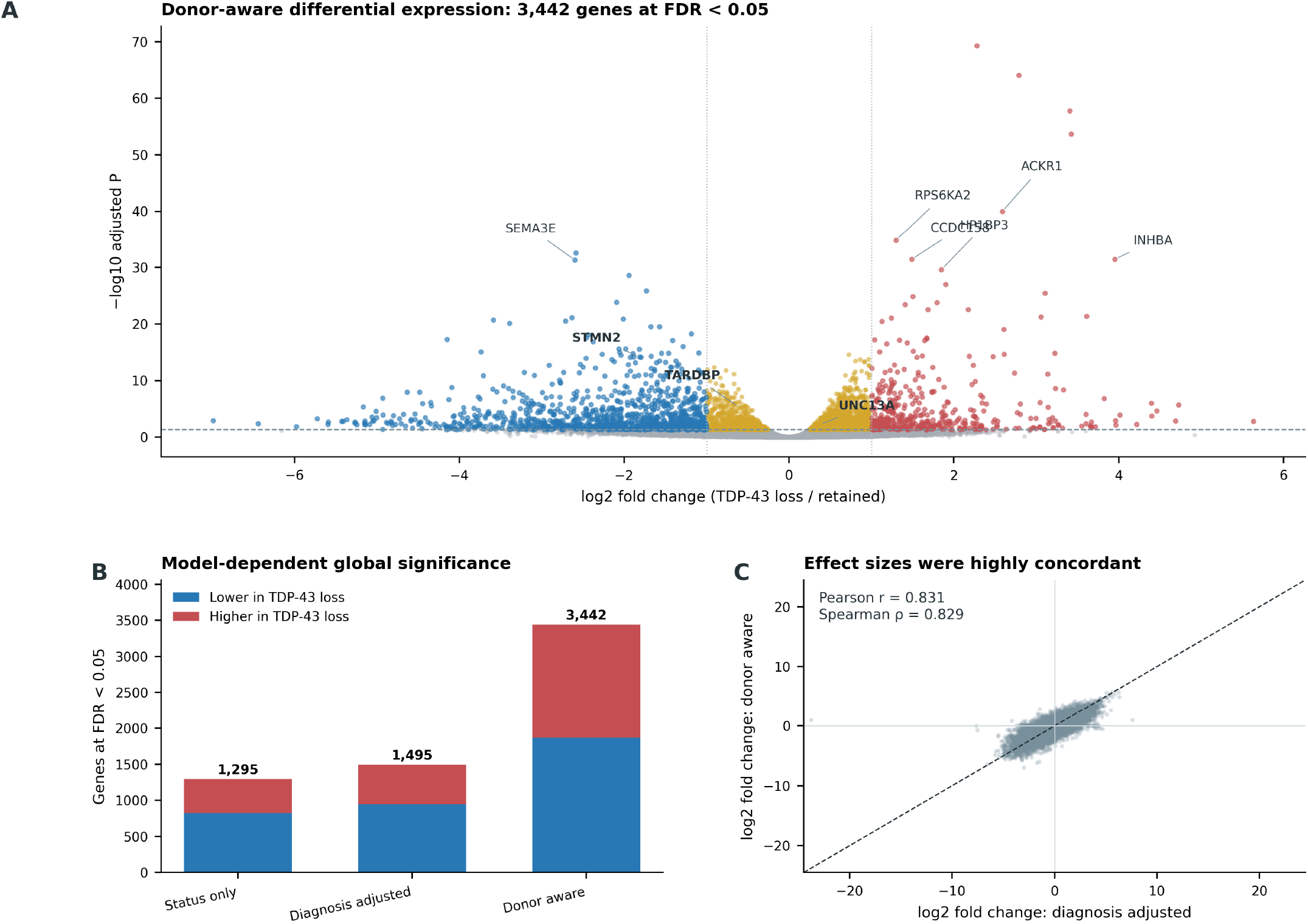
Differential expression and model sensitivity. (A) Donor-aware volcano plot for TDP-43 loss minus retained. Blue and red denote genes with FDR < 0.05 and absolute log2 fold change ≥ 1 in the lower and higher directions, respectively; gold denotes other FDR-significant genes. Selected protein-coding leaders and prespecified markers are labeled. (B) Numbers of FDR-significant genes in each model, partitioned by direction. (C) Gene-wise effect-size concordance between diagnosis-adjusted and donor-aware models; the dashed line is the identity line.

**Figure 4.**
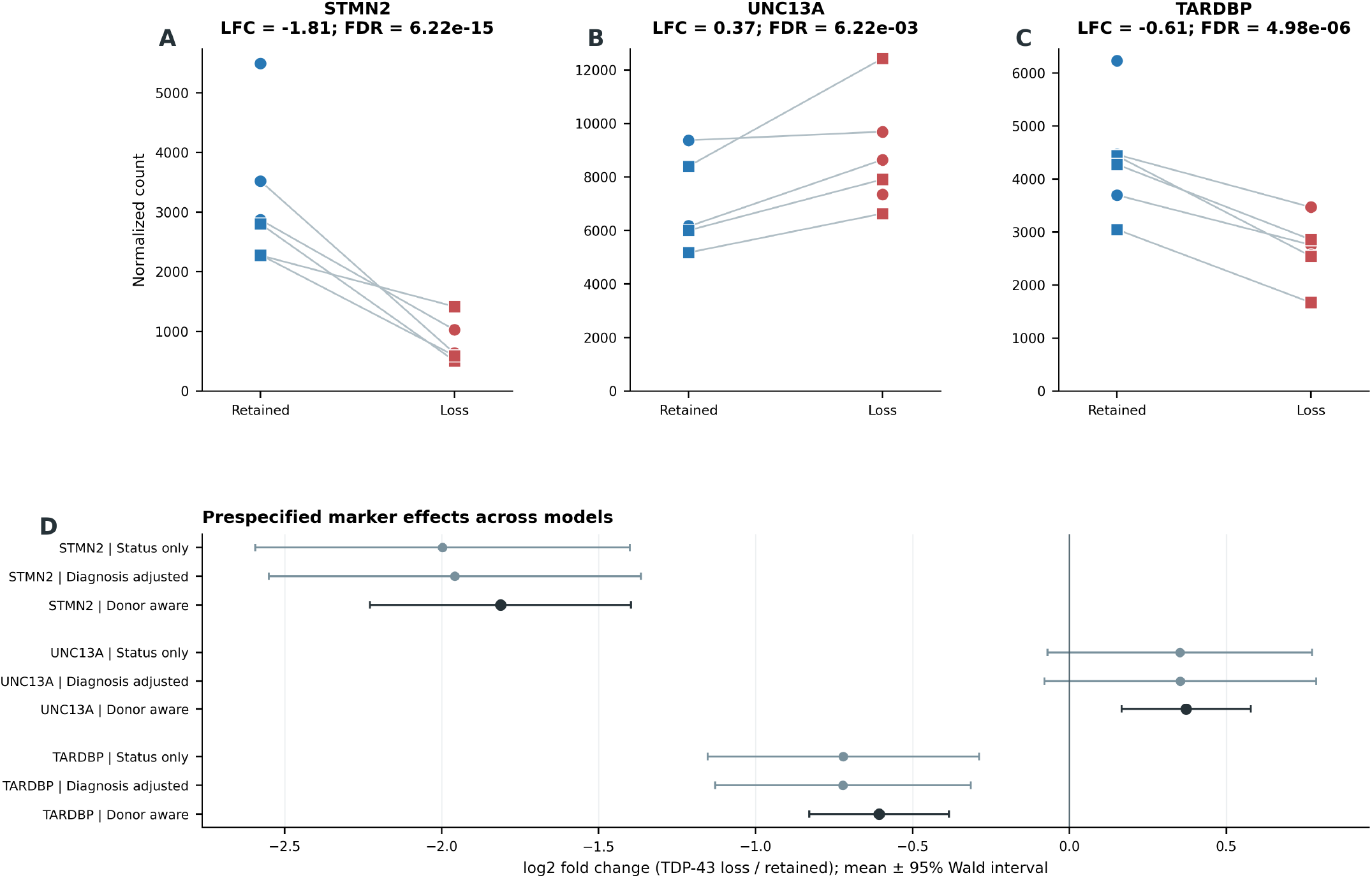
Prespecified marker expression across nuclear states and models. (A–C) Median-of-ratios-normalized counts for STMN2, UNC13A, and TARDBP. Lines connect complete matched donor pairs; shapes indicate diagnosis. Titles report donor-aware log2 fold change and FDR. (D) Marker log2 fold changes and 95% Wald intervals across status-only, diagnosis-adjusted, and donor-aware models. Negative values indicate lower expression in TDP-43-loss nuclei.

Despite the different numbers of discoveries, effect sizes from the donor-aware and diagnosis-adjusted models were strongly correlated across genes with finite estimates in both models (Pearson r = 0.831; Spearman ρ = 0.829; Fig. 3C; Table S7). Thus, broad effect-size direction was often conserved even when uncertainty and significance classifications differed. Volcano plots for the two sensitivity models are provided in Figure S2.

### Leading differential genes illustrate heterogeneity without establishing pathways

The 30 leading protein-coding genes showed coordinated groups of higher and lower expression across libraries, but the magnitude and pattern varied among donors (Fig. 5). Leading genes with lower expression in TDP-43-loss nuclei included SEMA3E, SST, NOS1, RPH3A, and SULF1; genes with higher expression included INHBA, RPS6KA2, HP1BP3, and SERPINE2. These genes are presented as descriptive leading features, not as evidence for a named pathway, cell-state transition, or therapeutic mechanism. The heatmap is based on the same data used for differential expression and therefore does not constitute external validation.

**Figure 5.**
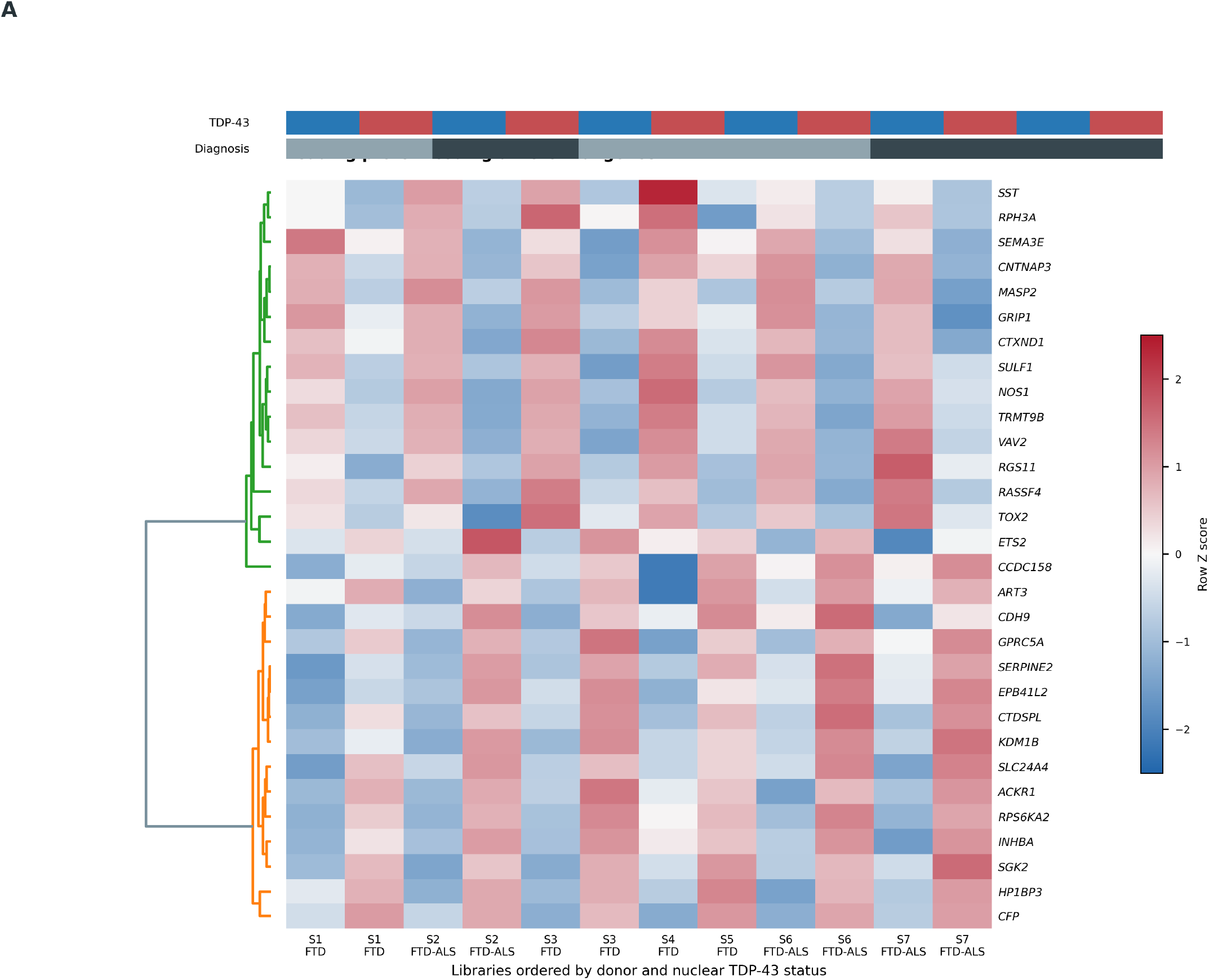
Leading protein-coding differential genes. The heatmap shows row Z scores of log2(normalized count + 1) for 30 protein-coding genes selected by adjusted P value among genes meeting FDR < 0.05, absolute log2 fold change ≥ 1, and baseMean ≥ 100. Rows use average-linkage clustering with correlation distance; columns remain ordered by donor and nuclear status. Top bars indicate nuclear TDP-43 status and diagnosis. This within-cohort display is descriptive, not independent validation.

### STMN2 depletion was robust across matched pairs and models

STMN2 showed a large, precisely estimated reduction in the donor-aware model: log2 fold change = −1.813, standard error = 0.212, 95% Wald interval −2.229 to −1.396, nominal P = 1.39 × 10^−17^, and FDR = 6.22 × 10^−15^ (Table 3; Fig. 4A,D). This effect corresponds to approximately 71.5% lower expression in TDP-43-loss nuclei. The association remained strong in the status-only model (log2 fold change = −1.997; FDR = 4.16 × 10^−8^) and diagnosis-adjusted model (log2 fold change = −1.958; FDR = 5.05 × 10^−8^).

**Table 3.** Donor-aware estimates for the three prespecified marker genes.

| Gene | baseMean | LFC (SE) | 95% Wald interval | FDR | Approx. change in loss |
|---|---|---|---|---|---|
| STMN2 | 2,005.98 | −1.813 (0.212) | −2.229 to −1.396 | 6.22 × 10<sup>−15</sup> | 71.5% lower |
| UNC13A | 7,830.44 | 0.372 (0.105) | 0.167 to 0.577 | 0.00622 | 29.4% higher |
| TARDBP | 3,503.54 | −0.606 (0.113) | −0.829 to −0.384 | 4.98 × 10<sup>−6</sup> | 34.3% lower |

LFC, log2 fold change for nuclear TDP-43 loss minus retained; SE, standard error. Approximate percent change is 100 × (2^LFC − 1). Cross-model estimates and intervals are provided in Table S4.

Normalized STMN2 counts were lower in the TDP-43-loss library for every complete donor pair (Fig. 4A; Table S8). The median within-pair descriptive log2 ratio was −1.942. Agreement across all five pairs, all three models, and the count-level visualization makes STMN2 the most robust biological finding of this reanalysis. Nevertheless, these data do not identify whether the reduction arose from premature polyadenylation, altered transcription, RNA stability, or another process.

### TARDBP was lower, whereas UNC13A whole-gene expression was not depleted

TARDBP, which encodes TDP-43, was moderately lower in TDP-43-loss nuclei in the donor-aware model (log2 fold change = −0.606; FDR = 4.98 × 10^−6^), with similar estimates in the sensitivity models (Fig. 4C,D). This gene-level association is compatible with the source nuclear-status annotation but does not measure nuclear protein abundance or distinguish transcriptional feedback from RNA-localization effects.

UNC13A showed the opposite whole-gene direction from a simple depletion expectation. Its donor-aware estimate was log2 fold change = 0.372 (95% Wald interval 0.167–0.577; FDR = 0.00622), equivalent to approximately 29.4% higher expression in TDP-43-loss nuclei (Fig. 4B,D). In contrast, UNC13A was not significant in the status-only or diagnosis-adjusted models (both FDR ≈ 0.50), indicating sensitivity to model specification. All five complete pairs had higher normalized UNC13A counts in the loss library, although the increase was very small in one pair.

This result should not be rewritten as UNC13A depletion, nor should it be interpreted as evidence against TDP-43-dependent UNC13A cryptic splicing. Whole-gene counts aggregate signal across constitutive and aberrant transcript species and reflect nuclear RNA rather than functional cytoplasmic protein. The present matrix cannot quantify UNC13A cryptic-exon inclusion, nonsense-mediated decay, or productive full-length UNC13A output.

## Discussion

This donor-aware secondary analysis supports three principal conclusions. First, nuclear TDP-43 loss is associated with extensive transcriptomic differences in FACS-isolated frontal-cortex neuronal nuclei, but the number of significant genes depends strongly on the design model. Second, STMN2 depletion is a robust within-cohort association, reproducible across all five matched pairs and all three statistical models. Third, UNC13A whole-gene expression was modestly higher only in the donor-aware model, demonstrating why whole-gene expression and cryptic-exon biology must be reported as separate layers of evidence.

The STMN2 finding is directionally consistent with experimental work identifying stathmin-2 as a TDP-43-sensitive mediator of neuronal growth and repair [3,4]. Its large effect size, narrow interval relative to the effect, and uniform direction across matched pairs strengthen the descriptive association. The result is also biologically coherent with loss of nuclear TDP-43 function rather than with nonspecific differences between unrelated disease groups. However, the present data are compatible with—not proof of—the established premature-polyadenylation mechanism. Demonstrating that mechanism would require event-level RNA evidence and, ideally, stathmin-2 protein measurement.

UNC13A illustrates the opposite risk: forcing gene-level output into a mechanistic narrative. Prior studies established TDP-43-sensitive UNC13A mis-splicing and depletion of functional UNC13A through cryptic-exon-related mechanisms [5,6]. The higher whole-gene estimate observed here could reflect nuclear transcript accumulation, donor-specific regulation, altered RNA stability, or model sensitivity, but these explanations remain speculative. More aberrant or unstable transcript species could contribute to total counts without increasing functional protein; conversely, a whole-gene increase does not demonstrate cryptic-exon inclusion. Only junction- and isoform-resolved analyses can adjudicate these possibilities.

The global results similarly caution against equating statistical yield with biological certainty. Donor-aware modeling changed both effect precision and the number of discoveries, while broad effect sizes remained concordant with the diagnosis-adjusted model. This pattern is plausible in a small, heterogeneous postmortem cohort where donor terms can absorb stable interindividual variation. It does not prove that every donor-aware discovery is causal or specific to TDP-43 loss. Leading-gene heatmaps are useful for inspecting coherent variation but cannot independently validate the model or establish pathway activation. We therefore avoided enrichment-based and mechanism-specific claims that were not directly tested.

The analysis also clarifies the appropriate next experiment. Reprocessing all 14 eligible sorted libraries from FASTQ files with a recorded genome build and annotation release would remove the present subset-selection ambiguity and enable uniform alignment. Junction-aware quantification, supplemented by intron-retention and alternative-polyadenylation analyses, could directly test STMN2 and UNC13A processing. Long-read RNA sequencing could connect distant splice and polyadenylation events within the same molecules. Targeted RT–PCR across the relevant events, quantitative protein assays, and independent cohorts would then be required to establish functional consequences and generalizability.

## Limitations

Several limitations constrain interpretation. The cohort is small: 12 libraries from seven donors, with only five complete matched pairs. Two donors contribute a single nuclear state, and two eligible sorted libraries from the full source series are absent from the supplied matrix for undocumented reasons. This creates potential selection bias and prevents a complete 14-library sensitivity analysis from the available matrix.

The source cohort contains FTD and FTD–ALS postmortem tissue without healthy controls, and source metadata annotate a C9orf72 repeat-expansion background. Findings therefore describe associations within diseased donors and cannot distinguish disease onset from progression, establish effects in sporadic ALS/FTD, or support comparisons with healthy aging. Diagnosis and sex distributions are also too limited for reliable subgroup-specific estimates. Postmortem interval, agonal state, and other technical covariates were not available for expanded adjustment.

The measurements are restricted to NeuN-positive neuronal nuclei from frontal cortex. They do not represent unsorted bulk cortex, glia, cytoplasmic RNA, other brain regions, or protein abundance. Nuclear TDP-43 status was inherited from the source annotation and was not independently verified by new imaging or biochemical assays. Sorting purity and per-nucleus protein measurements cannot be reassessed from the count matrix.

The analysis is gene-level. It provides no direct evidence of cryptic-exon inclusion, PSI, splice-junction usage, isoform abundance, nonsense-mediated decay, or cryptic peptides. In particular, neither STMN2 premature polyadenylation nor UNC13A cryptic splicing was measured. Whole-gene UNC13A expression must not be used as a surrogate for functional UNC13A protein.

Finally, the original FASTQ processing was not rerun. Although the supplied matrix exactly matches featureCounts Assigned counts, the original genome build, annotation release, alignment settings, and complete quality-control trail could not be reconstructed. Annotation through MyGene.info may change over time and some identifiers remain unmapped. All discoveries derive from the same cohort, with no external validation; leading-gene displays and model comparisons are exploratory and should guide confirmation rather than serve as clinical evidence.

## Conclusions

Donor-aware reanalysis of FACS-isolated neuronal nuclei identifies robust STMN2 depletion associated with nuclear TDP-43 loss in this FTD/FTD–ALS cohort. The association is supported by all five matched donor pairs and by sensitivity analyses, making it a strong candidate for targeted mechanistic follow-up. Global differential expression and UNC13A whole-gene estimates are more model-dependent. Cryptic-exon inclusion and downstream RNA or protein consequences remain unmeasured and must be tested with junction-resolved and experimental approaches before mechanistic conclusions are drawn.

## Declarations

### Ethics statement

This work used only publicly available, deidentified secondary data and involved no new recruitment, specimen collection, or interaction with human participants. Ethical approval and consent for the original study were the responsibility of the primary investigators [8].

### Data availability

The source data are available through GEO series GSE126542 and BioProject PRJNA522295. The exact 12-library count matrix, featureCounts summary, verified metadata, differential-expression results, derived tables, and figure-generation outputs analyzed here are included in the accompanying preprint package.

### Code availability

All scripts used for validation, differential expression, annotation, and figure generation are included in the accompanying code directory and are available through the project GitHub repository: https://github.com/abdullahchaudhary360/Tdp43-Neuronal-Nuclei-Reanalysis. A configurable FASTQ-to-counts workflow is also supplied but was not executed for the reported results.

### Author contributions

Abdullah Chaudhary: conceptualization, project leadership and administration, methodology, software and pipeline development, quality-control and trimming work, formal analysis, validation, visualization, and writing—original draft, review, and editing. Abdullah Chaudhary conceived and led the project, wrote the analysis code, designed and set up the analyses, and conducted and coordinated the main project work. Saiqi Yao: data gathering and curation, DESeq2 analysis support, generation of selected figures, and manuscript writing support. Both authors contributed collaboratively to the project and approved the final manuscript.

## Acknowledgments

The authors thank Liu et al. for releasing the GSE126542 data and metadata. Sanbiomics tutorials are acknowledged as general training resources for pipeline development; no tutorial output was used as study data or as evidence in this reanalysis.

## Generative-AI assistance disclosure

During manuscript preparation, a generative-AI assistant (Kimi, Moonshot AI) assisted with file auditing, code organization, figure formatting, and language editing. The analysis outputs were generated from the supplied data and scripts, and the authors remain responsible for verifying the analyses, references, interpretations, and final manuscript. No AI system is listed as an author.

## Supplementary figures

**Figure S1.**
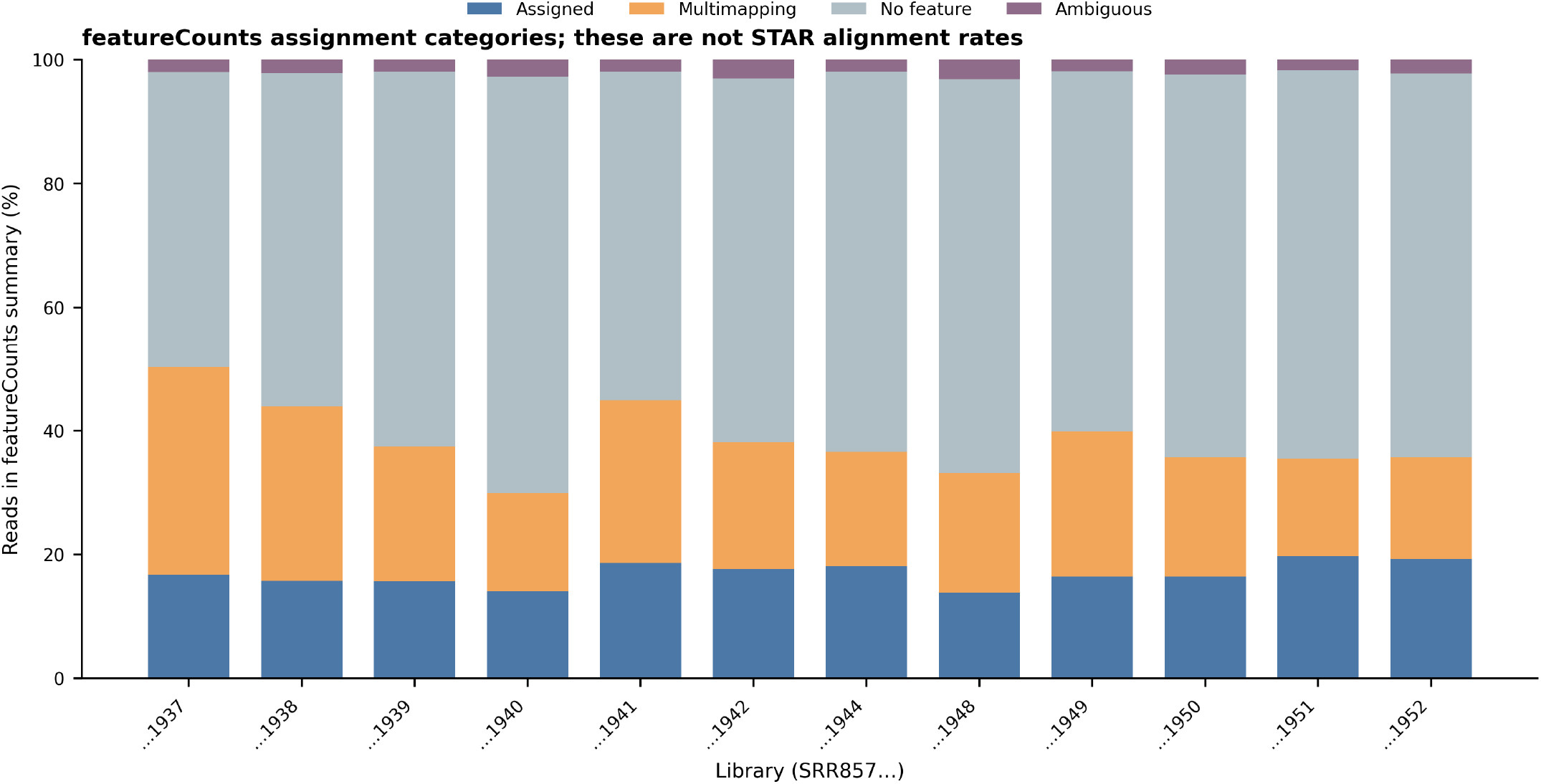
featureCounts assignment categories. Stacked bars show Assigned, Unassigned_MultiMapping, Unassigned_NoFeatures, and Unassigned_Ambiguity reads as percentages of the reads represented in the featureCounts summary. These categories are not STAR alignment rates.

**Figure S2.**
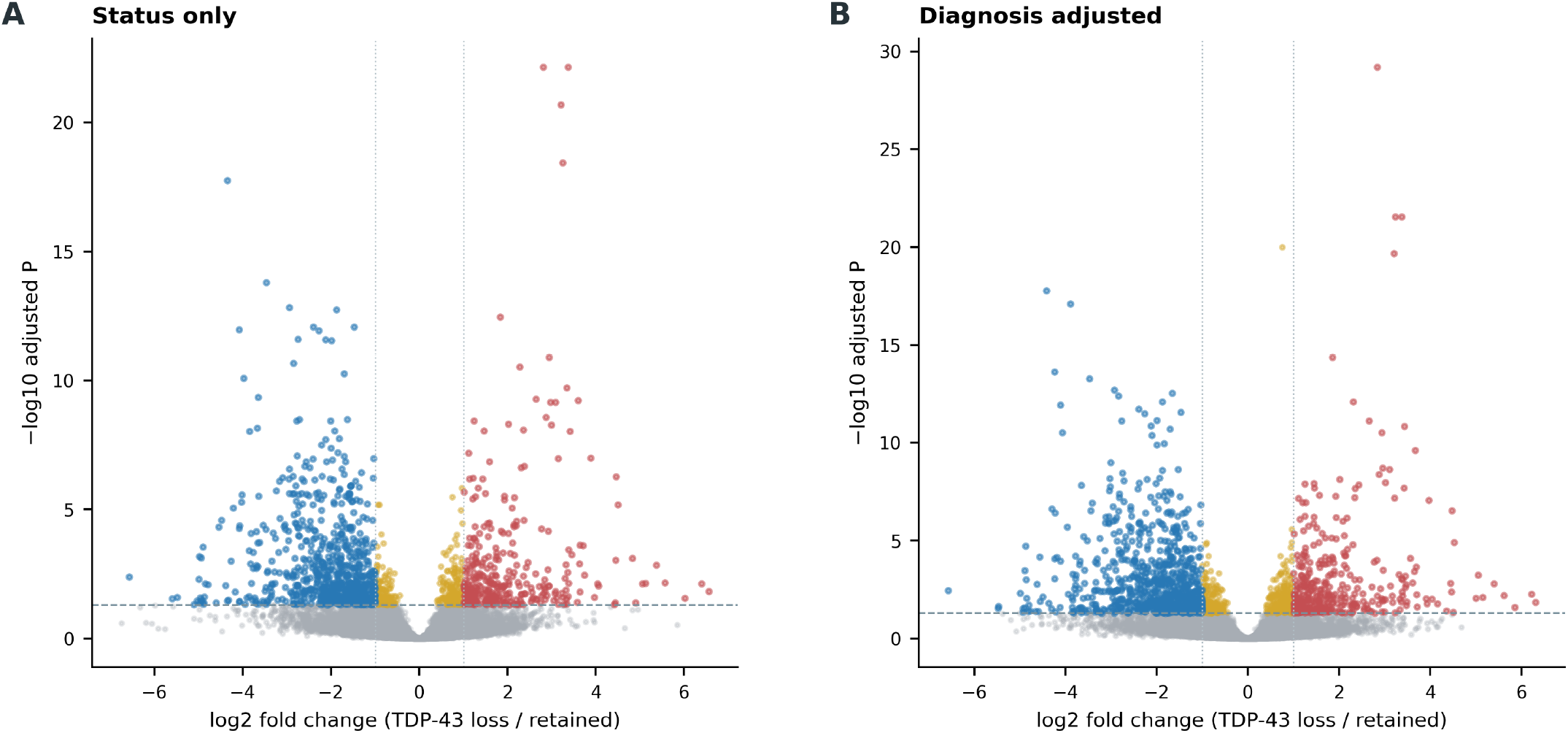
Sensitivity-model differential expression. Volcano plots for (A) ∼ nuclear_status and (B) ∼ diagnosis + nuclear_status. Thresholds and colors match Figure 3A; all contrasts are nuclear TDP-43 loss minus retained.

**Figure S3.**
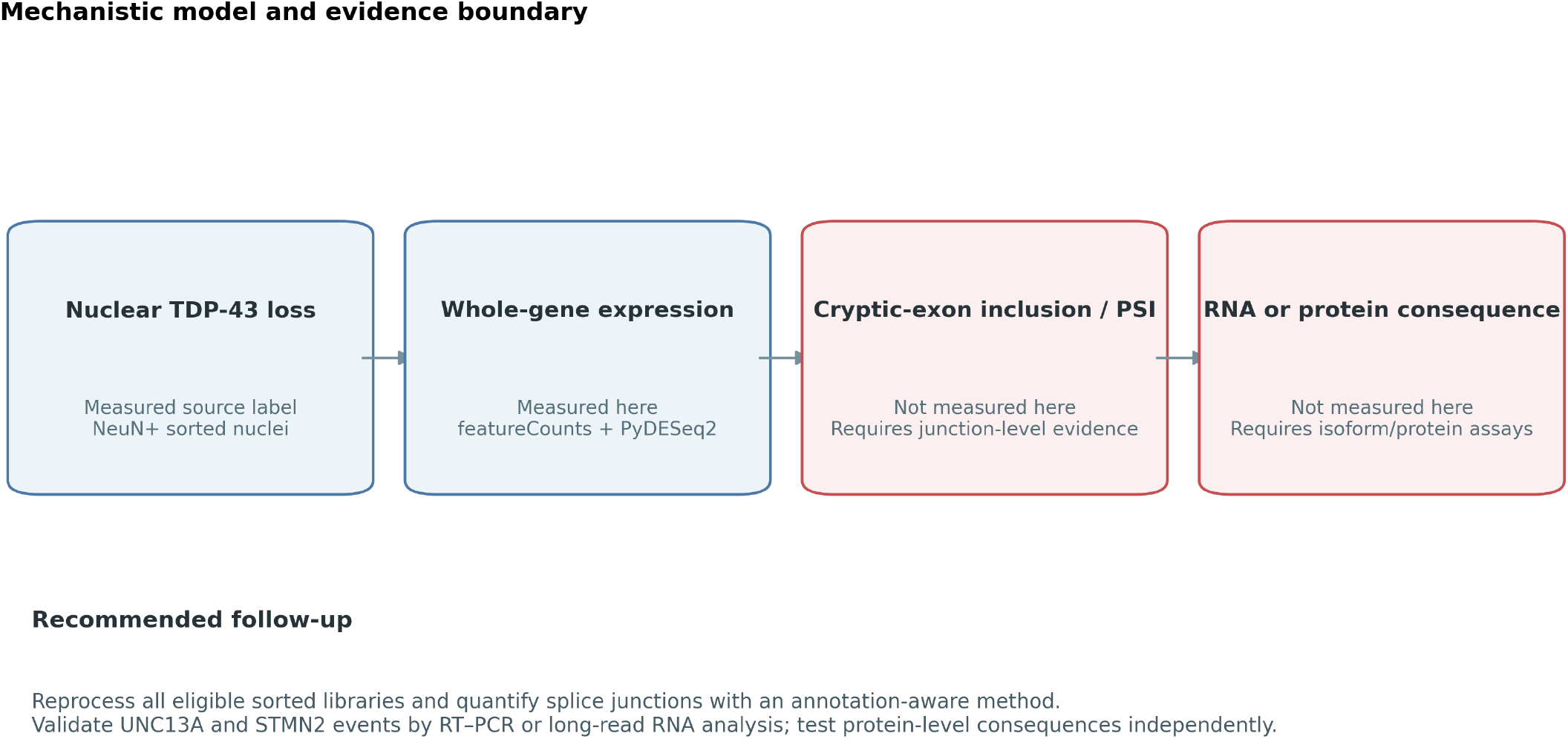
Mechanistic model and evidence boundary. Source nuclear-status labels and whole-gene expression were measured or available to this reanalysis. Cryptic-exon inclusion/PSI and downstream RNA or protein consequences were not measured and require junction-level, isoform-resolved, or experimental assays. Recommended follow-up steps are shown.

## Supplementary tables

The following editable tab-delimited files accompany the manuscript.

- **Table S1** — Verified sample-level metadata for the 12 analyzed libraries. File: Table_S1_sample_metadata.tsv
- **Table S2** — Annotated donor-aware genes with FDR < 0.05. File: Table_S2_donor_aware_significant_genes.tsv
- **Table S3** — Fifty leading donor-aware genes by adjusted P value. File: Table_S3_top_genes.tsv
- **Table S4** — Marker estimates, standard errors, Wald intervals, and FDRs for all three models. File: Table_S4_marker_model_statistics.tsv
- **Table S5** — Global and marker-level summaries across models. File: Table_S5_model_comparison.tsv
- **Table S6** — PCA coordinates and sample annotations. File: Table_S6_PCA_coordinates.tsv
- **Table S7** — Gene-wise donor-aware and diagnosis-adjusted effect sizes. File: Table_S7_effect_size_concordance.tsv
- **Table S8** — Normalized STMN2, UNC13A, and TARDBP counts by library. File: Table_S8_marker_normalized_counts.tsv
- **Table S9** — Selection statistics for the 30 heatmap genes. File: Table_S9_heatmap_gene_selection.tsv
- **Table S10** — Descriptive median-of-ratios size factors. File: Table_S10_size_factors.tsv

