## Supplementary Figures S1-S3 for "Donor-Aware Reanalysis of Neuronal Nuclei Supports STMN2 Depletion Associated with Nuclear TDP-43 Loss in FTD and ALS": Figure_S1_featureCounts_assignment.pdf

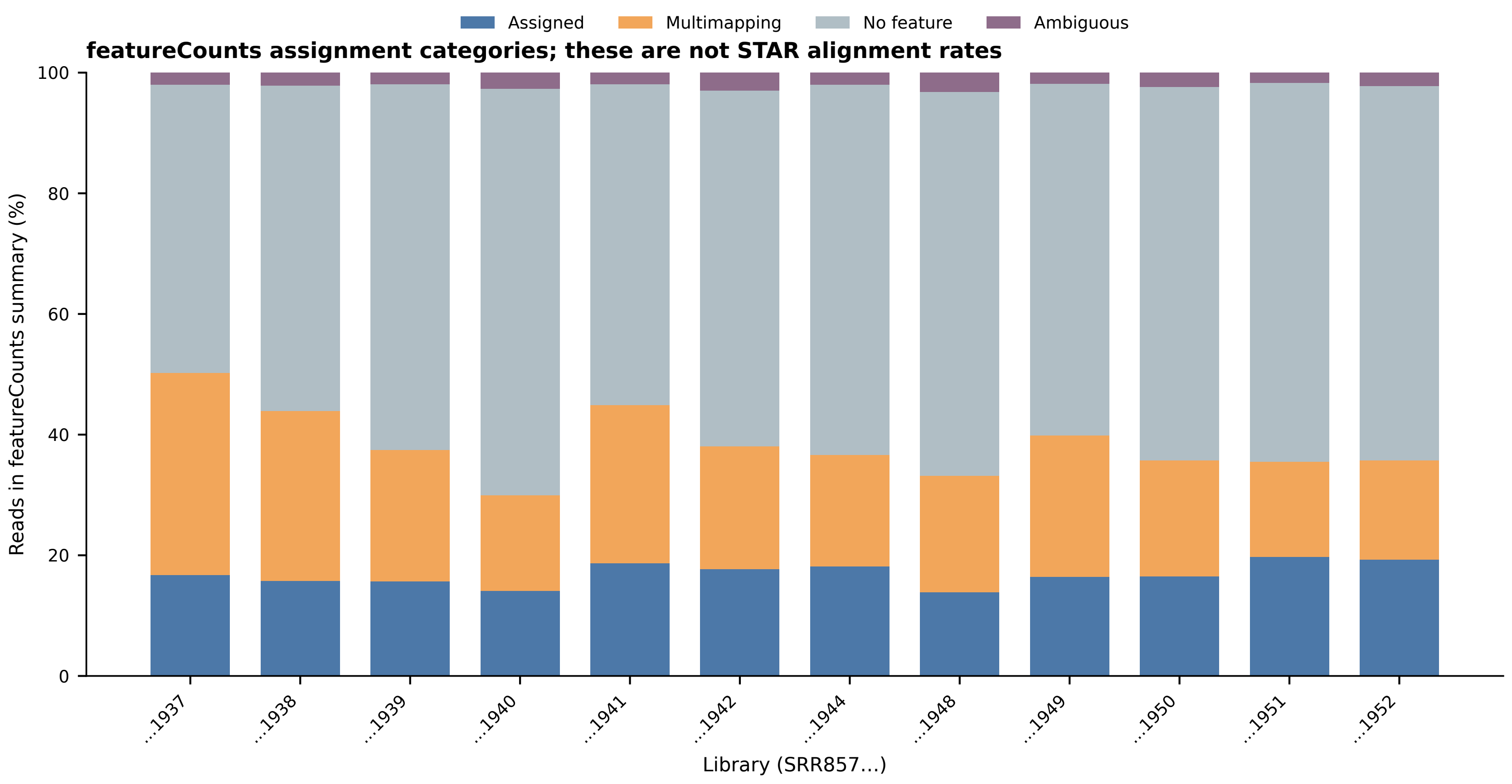
Assigned
Multimapping
No feature
Ambiguous
featureCounts assignment categories; these are not STAR alignment rates
Reads in featureCounts summary (%)
0
20
40
60
80
100
…1937
…1938
…1939
…1940
…1941
…1942
…1944
…1948
…1949
…1950
…1951
…1952
Library (SRR857…)
