## Supplementary Figures S1-S3 for "Donor-Aware Reanalysis of Neuronal Nuclei Supports STMN2 Depletion Associated with Nuclear TDP-43 Loss in FTD and ALS": Figure_S3_cryptic_exon_evidence_boundary.pdf

### Mechanistic model and evidence boundary

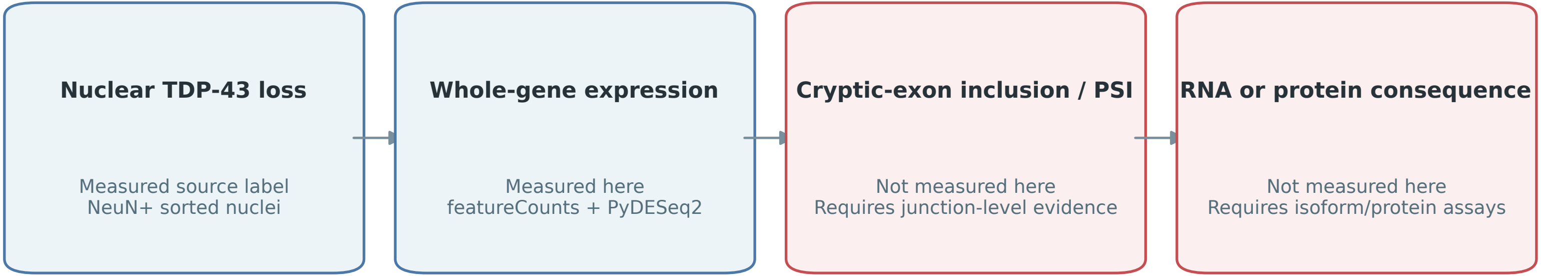


#### Recommended follow-up

Reprocess all eligible sorted libraries and quantify splice junctions with an annotation-aware method.
Validate UNC13A and STMN2 events by RT–PCR or long-read RNA analysis; test protein-level consequences independently.
