## Supplementary figures and images for "Donor-Aware Reanalysis of Neuronal Nuclei Supports STMN2 Depletion Associated with Nuclear TDP-43 Loss in FTD and ALS"

### Figure_S2_sensitivity_volcanoes.pdf

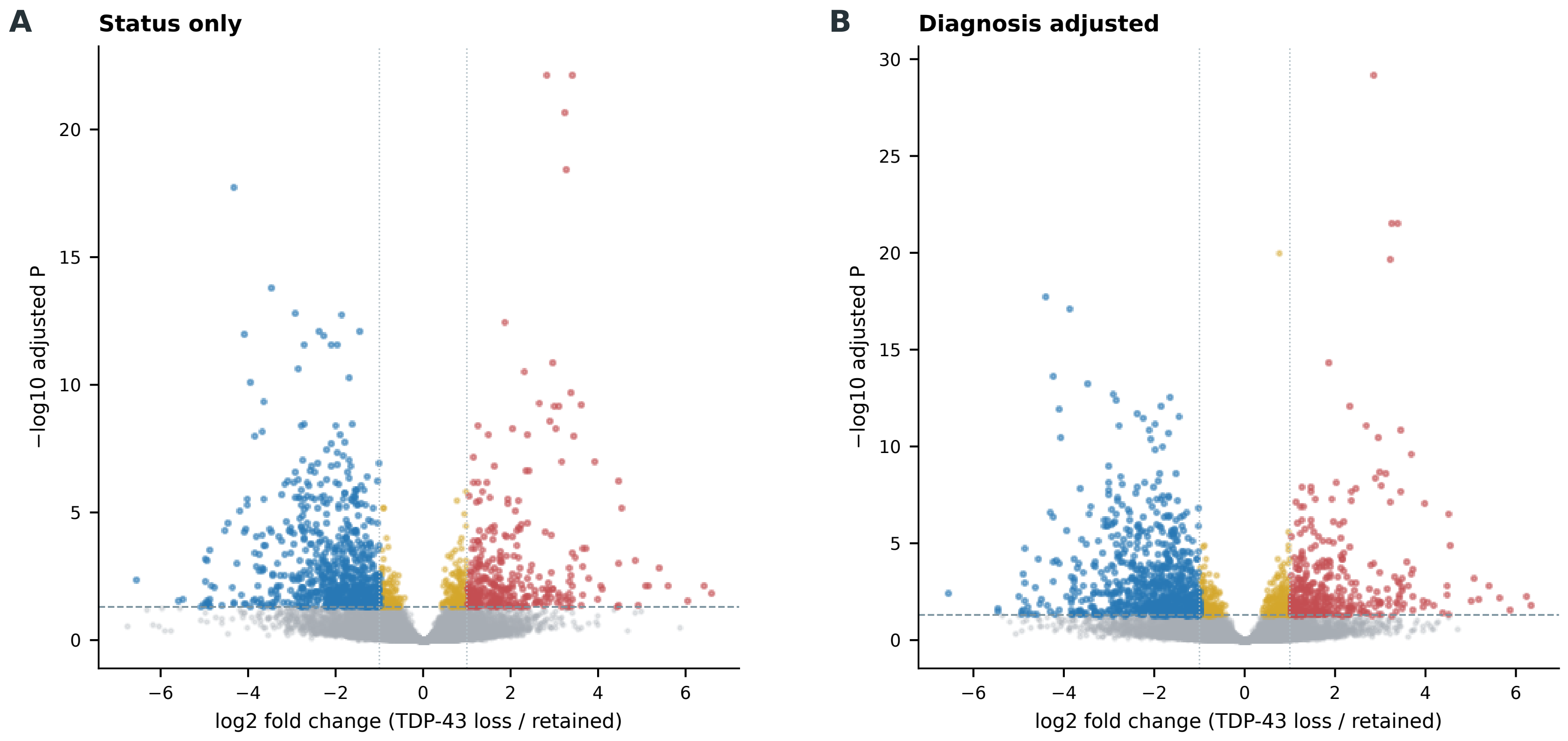
A
Status only
B
Diagnosis adjusted
−log10 adjusted P
log2 fold change (TDP-43 loss / retained)
0
5
10
15
20
25
30
−6
−4
−2
0
2
4
6
